# Signaling incentive and drive in the primate ventral pallidum for motivational control of goal-directed action

**DOI:** 10.1101/420851

**Authors:** Atsushi Fujimoto, Yukiko Hori, Yuji Nagai, Erika Kikuchi, Kei Oyama, Tetsuya Suhara, Takafumi Minamimoto

## Abstract

Processing incentive and drive is essential for control of goal-directed behavior. The limbic part of the basal ganglia has been emphasized in these processes, yet the exact neuronal mechanism has remained elusive. In this study, we examined the neuronal activity of the ventral pallidum (VP) and its upstream area, the rostromedial caudate (rmCD), while two male macaque monkeys performed an instrumental lever-release task, in which a visual cue indicated the forthcoming reward size. We found that the activity of some neurons in VP and rmCD reflected the expected reward-size transiently following the cue. Reward-size coding appeared earlier and stronger in VP than in rmCD. We also found that the activity in these areas was modulated by the satiation level of monkeys, which also occurred more frequently in VP than in rmCD. The information regarding reward-size and satiation-level was independently signaled in the neuronal populations of these areas. The data thus highlighted the neuronal coding of key variables for goal-directed behavior in VP. Furthermore, pharmacological inactivation of VP induced more severe deficit of goal-directed behavior than inactivation of rmCD, which was indicated by abnormal error repetition and diminished satiation effect on the performance. These results suggest that VP encodes incentive value and internal drive, and plays a pivotal role in the control of motivation to promote goal-directed behavior.

**Significance Statement:** The limbic part of the basal ganglia has been emphasized in the motivational control of goal-directed action. Here, we investigated how the ventral pallidum (VP) and the rostromedial caudate (rmCD) encode incentive value and internal drive, and control goal-directed behavior. Neuronal recording and subsequent pharmacological inactivation revealed that VP had stronger coding of reward size and satiation level than rmCD. Reward size and satiation level were independently encoded in the neuronal population of these areas. Furthermore, VP inactivation impaired goal-directed behavior more severely than rmCD inactivation. These results highlighted the central role of VP in the motivational control of goal-directed action.

## Introduction

Motivational control over the purposeful action, or goal-directed behavior, is essential for gaining reward from an environment through knowledge about the association between action and its consequence (Dickinson, 1985; Dickinson and Balleine, 1994). Impairment of motivational control of goal-directed behavior promotes autonomic (habitual) control of actions, and this is evident in addictive disorders (Everitt and Robbins, 2005; Ersche et al., 2016). Goal-directed behavior is regulated by two factors — the incentive value of the goal (reward) and the internal drive (physiological state) of an agent (Berridge, 2004; Zhang et al., 2009). Accordingly, motivational processes would be governed by the signals related to these two factors (i.e., motivational value), although their neural mechanism has remained largely unknown.

The ventral pallidum (VP), an output nucleus of ventral basal ganglia, is posited in the heart of the limbic system (Haber et al., 1985; Groenewegen et al., 1993; Ray and Price, 1993; Mai and Paxinos, 2011) and has been strongly implicated in reward processing (Smith et al., 2009; Castro et al., 2015; Root et al., 2015). Neuronal activity in VP has been shown to reflect the incentive value of reward cue in rodents (Tindell et al., 2004; Ahrens et al., 2016) and monkeys (Tachibana and Hikosaka, 2012; Saga et al., 2017). Pharmacological manipulation of VP disrupted normal reward-based behavior in monkeys (Tachibana and Hikosaka, 2012; Saga et al., 2017). Dysregulation of its neuronal activity induces addiction-like behavior in mice (Mahler et al., 2014; Faget et al., 2018). Collectively, these results suggest a significant contribution of VP to goal-directed behavior.

Other studies have also focused on the rostromedial part of the caudate nucleus (rmCD), one of the upstream structures of VP (Haber et al., 1990), as making a significant contribution to goal-directed behavior. The rmCD receives projections from the lateral orbitofrontal cortex (OFC) (Haber and Knutson, 2010; Averbeck et al., 2014), and attenuation of OFC-striatal activity promotes habitual control of action over the goal-directed action in rodents (Yin et al., 2005; Gremel and Costa, 2013; Gremel et al., 2016). In monkeys, neuronal activity in the middle caudate including rmCD reflected reward size (Nakamura et al., 2012), and silencing of rmCD neurons induced a loss of reward-size sensitivity and disrupted goal-directed performance (Nagai et al., 2016).

Considering the direct anatomical connection from rmCD to VP, although they may interplay and contribute to a goal-directed control of action, the exact mechanism still remains unclear. One possible mechanism is that rmCD relays the signal derived from the OFC, which reflects the incentive value and internal drive (Rolls, 2006), and is further strengthened through the convergence projection to the VP (Haber and Knutson, 2010; Averbeck et al., 2014). Other mechanisms are also conceivable, as, for example, that incentive value and internal drive are processed outside the striatum, such as in the amygdala and hypothalamus (Burton et al., 1976; Paton et al., 2006), which project to VP to compute motivational value through its downstream structures. To clarify the neural mechanism of motivational control of action, therefore, a comparison of the neuronal coding of VP and rmCD in terms of incentive and drive is essential.

In this study, we analyzed the single-unit activities of these two areas while macaque monkeys performed an instrumental lever-release task, in which a visual cue indicated the forthcoming reward size (Minamimoto et al., 2009). As this task design permits us to infer the impact of incentive value (i.e., reward size) and internal drive (i.e., satiation level of monkeys) on performance, we assessed the neuronal correlate of the two factors, and compared neuronal coding between the two areas. With a population-level comparison, we found that the coding of reward size and satiation level in VP was greater than that in rmCD. Pharmacological inactivation of VP further examined the causal contribution of the neuronal activity to goal-directed action. Our results suggest a central role of VP in motivational control of goal-directed behavior and may provide implication for the neural mechanism of addictive disorders.

## Materials and Methods

### Subjects

Four male rhesus monkeys (*Macaca mulatta*, 5.7-7.2 kg) were used in this study. Two were used for neuronal recording (monkeys TA and AP) and the other two for local inactivation experiments (monkeys RI and BI). Monkey RI was also used in the previous rmCD inactivation study (Nagai et al., 2016). All surgical and experimental procedures were approved by the Animal Care and Use Committee of the National Institutes for Quantum and Radiological Science and Technology and were in accordance with the guidelines published in the NIH Guide for the Care and Use of Laboratory Animals.

### Behavioral task

The monkeys squatted on a primate chair inside a dark, sound-attenuated, and electrically shielded room. A touch-sensitive lever was mounted on the chair. Visual stimuli were displayed on a computer video monitor in front of the animal. Behavioral control and data acquisition were performed using a real-time experimentation system (REX) (Hays Jr et al., 1982). Presentation software was used to display visual stimuli (Neurobehavioral Systems Inc., Berkeley, CA).

All four monkeys were trained to perform the reward-size task (Minamimoto et al., 2009) (Fig. 1a). In each of the trials, the monkey had the same requirement of obtaining one of four sizes of liquid rewards (1, 2, 4, or 8 drops, 1 drop = ca. 0.1 mL). A trial began when a monkey gripped a lever. A visual cue and a red spot appeared sequentially, with a 0.4 s interval, at the center of the monitor. After a variable interval (0.5 -1.5 s), the central spot turned to green (‘go’ signal), and the monkey had to release the lever within the reaction time window (0.2-1.0 s). If the monkey released the lever correctly, the spot turned to blue (0.2-0.4 s), and then a reward was delivered. The next trial began following an inter-trial interval (ITI, 1.5 s). When trials were performed incorrectly, they were terminated immediately (all visual stimuli disappeared), and the next trial began with the same reward condition following the ITI. There were two types of errors: premature lever releases (lever releases before or no later than 0.2 s after the appearance of the go signal, named “early errors”) and failures to release the lever within 1.0 s after the appearance of the go signal (named “late errors”).

**Figure 1.**
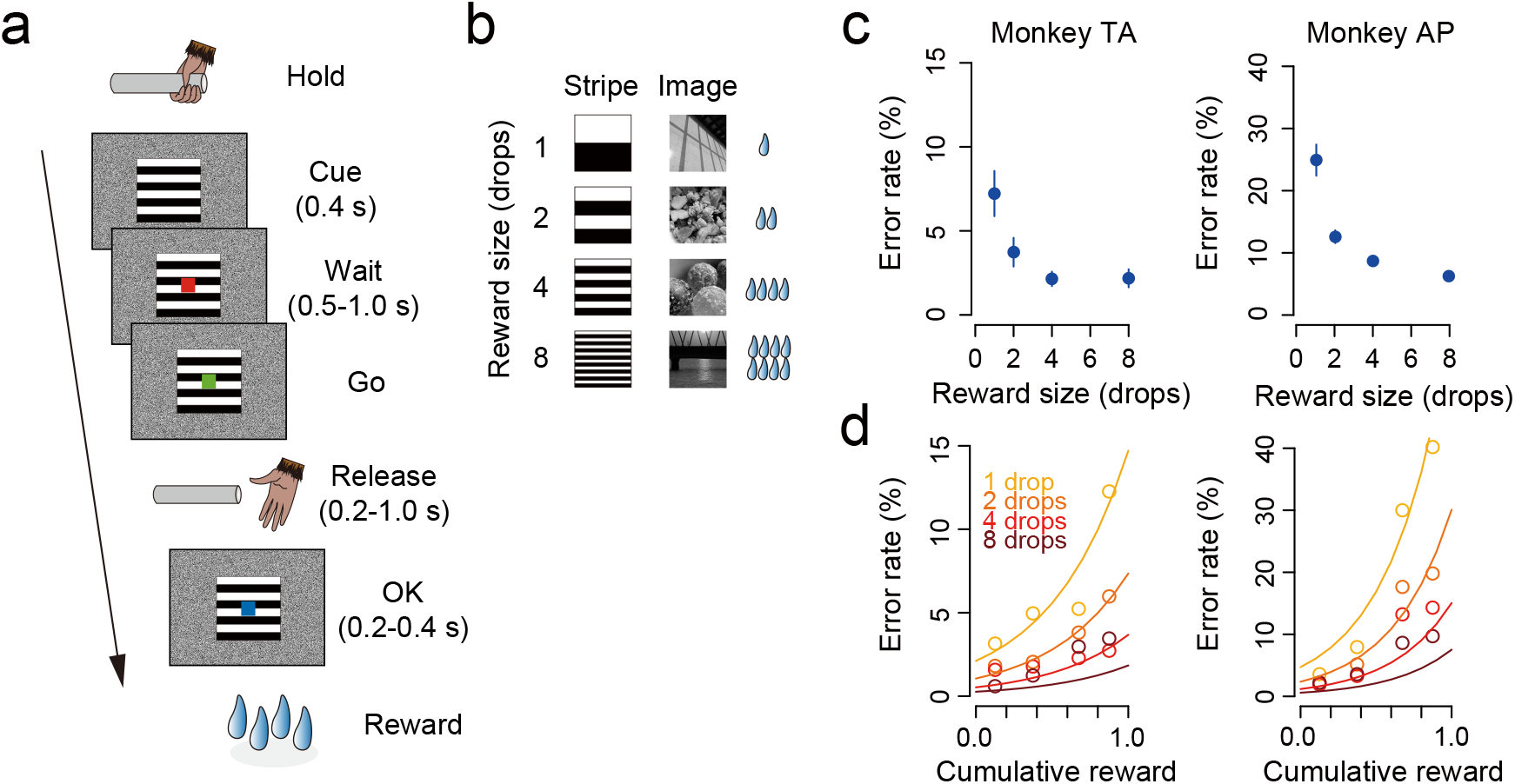
Reward-size task and behavioral performance. ***a***. Sequence of a trial. ***b***. Cue stimuli. Either stripe set (left column) or image set (right column) was used to inform the reward size (1, 2, 4, 8 drops of liquid). ***c***. Error rate (mean ± SEM) as a function of reward size for monkeys TA and AP, respectively. ***d***. Mean error rate as a function of normalized cumulative reward for the two monkeys. Each color indicates reward size. Curves were best fit of Eq. 1 and 2 with *c* = 2.1, *λ* = 1.9, for monkey TA; *c* = 4.7, *λ* = 2.5, for monkey AP.

The size of the reward was chosen randomly and was indicated by visual cues at the beginning of each trial. Two sets of cues were used: a stripe set (for monkeys TA, RI, and BI) and an image set (for monkey AP) (Fig. 1b). The monkeys used for electrophysiology (monkeys TA and AP) were trained with different cue sets so that we could interpret the reward-related neuronal signal irrespective of the visual features of the cue stimuli.

Prior to the experiment with the reward-size task, all monkeys had been trained to perform color discrimination trials in a cued multi-trial reward schedule task for more than 3 months.

### Surgery

After behavioral training, a surgical procedure was carried out to implant one or two recording chambers and a head fixation device under general isoflurane anesthesia (1-2%). The angles of the chamber(s) were vertical (monkeys TA, AP, and BI) or 20° tilted from the vertical line (monkey RI) in the coronal plane. Prior to surgery, overlay magnetic resonance (MR) and X-ray computed tomography (CT) images were created using PMOD image analysis software (PMOD Technologies Ltd, Zurich, Switzerland) to estimate the stereotaxic coordinates of the target brain structures. MR images at 7T (Bruker Corp., Billerica, MA) and CT images (3D Accuitomo170: J. Morita Corp., Osaka, Japan) were obtained under anesthesia (propofol 0.2-0.6 mg/kg/min, i.v.).

### Neuronal recordings

Single-unit activity was recorded from monkeys TA and AP while they performed the reward-size task. We analyzed all successfully isolated activities and held at least 10 trials for each reward condition. Action potentials of single neurons were recorded from VP and rmCD using a glass-coated 1.0 MΩ tungsten microelectrode (Alpha Omega Engineering Ltd., Nazareth, Israel). A guide tube was inserted through the grid hole in the implanted recording chamber into the brain, and the electrodes were advanced through the guide tube by means of a micromanipulator (MO-97A: Narishige Co., Ltd., Tokyo, Japan). Spike sorting to isolate single neuron discharges was performed with a time-window algorithm (TDT-RZ2: Tucker Davis Technologies Inc., Alachua, FL). The timing of action potentials was recorded together with all task events at millisecond precision.

For the VP recordings, we targeted the region just below the anterior commissure (AC) in the +0-1 mm coronal plane (Fig. 2a). VP neuron was characterized by high spontaneous firing rate with phasic discharge to the task events (Tachibana and Hikosaka, 2012). For the rmCD recordings, we targeted the area within 2-4 mm laterally and 2-7 mm ventrally from the medial and upper edge of the caudate nucleus, in the +4-5 mm coronal plane (Fig. 2b). The rmCD neurons were classified into three subtypes based on the electrophysiological criteria (Aosaki et al., 1995; Yamada et al., 2016). The presumed medium-spiny projection neurons (PANs: phasically-active neurons) were characterized by low spontaneous firing and phasic discharge to the task events, while the presumed cholinergic interneurons (TANs: tonically-active neurons) were characterized by broad spike width (valley-to-peak width) and tonic firing around 3.0-8.0 Hz. The presumed parvalbumin-containing GABAergic interneurons (FSNs: fast-spiking neurons) were characterized by narrow spike width and relatively higher spontaneous firing than other types of caudate neurons. A spike-width analysis was performed using the Off-line sorter (Plexon, Dallas, TX).

**Figure 2.**
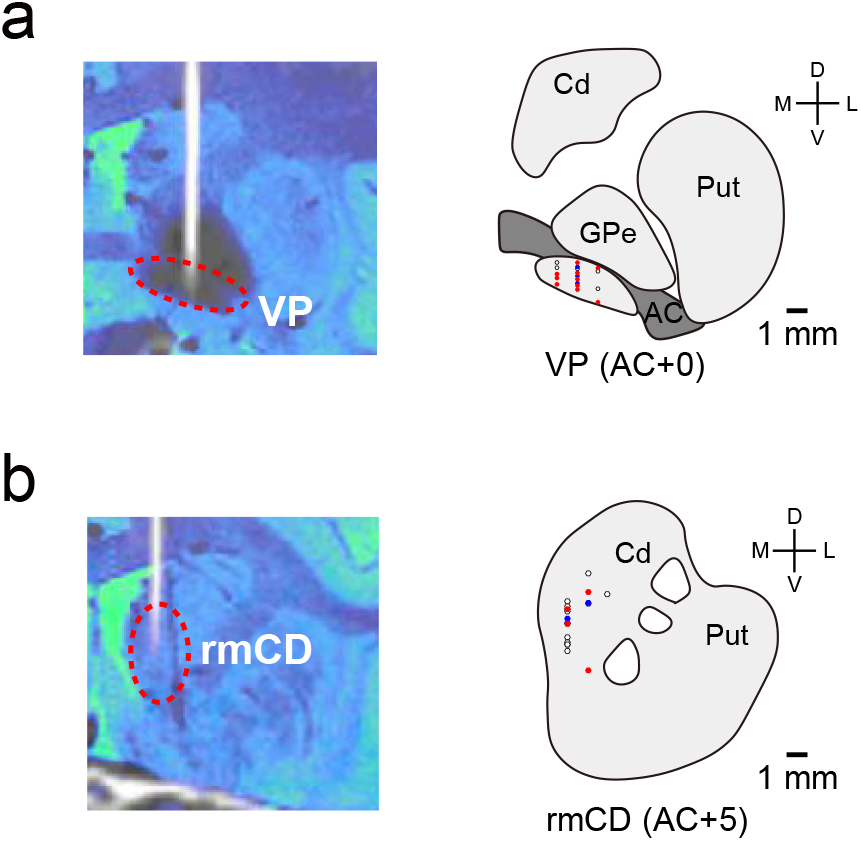
Recording sites in VP and rmCD. ***a-b***. Recording sites of VP and rmCD, respectively. Left: CT/MR fusion image showing the position of an electrode. Right: Schematic pictures representing the locations of the recorded neurons: positive reward-size coding neurons (red), negative reward-size coding neurons (blue), and non-coding neurons (white). Representative slices from monkey TA were used. Cd: Caudate nucleus, Put: Putamen, GPe: External segment of the globus pallidus, AC: Anterior commissure.

To reconstruct the recording location, electrodes were visualized using CT scans after each recording session, and the positions of the tip were mapped onto the MR image using PMOD.

### Muscimol microinjection

To achieve neuronal silencing, GABA_A_ agonist muscimol (Sigma-Aldrich Co., St. Louis, MO) was injected bilaterally into VP (monkeys RI and BI) using the same procedures as reported previously (Nagai et al., 2016). We used two stainless steel injection cannulae inserted into the caudate (O.D. 300 μm: Muromachi-Kikai Co. Ltd., Tokyo, Japan), one in each hemisphere. Each cannula was connected to a 10-μL microsyringe (#7105KH: Hamilton Company, Reno, NV) via polyethylene tubing. These cannulae were advanced through the guide tube by means of an oil-drive micromanipulator. Muscimol (3 μg/1 μL saline) was injected at a rate of 0.2 μL/min by auto-injector (Legato210: KD Scientific Inc., Holliston, MA) for a total volume of 2 μL in each side. The behavioral session (100 min) was started soon after the injection was finished. We performed at most one inactivation study per week. For a control, we performed sham experiments at other times, in which the time-course and mechanical settings were set identical to the muscimol session. At the end of each session, a CT image was obtained to visualize the injection cannulae in relation to the chambers and skull. The CT image was overlaid on an MR image by using PMOD to assist in identifying the injections sites.

### Experimental design and statistical analysis

All statistical analyses were performed with R Statistical Package. For the behavioral analysis, the data obtained from the two monkeys were analyzed. The dependent variables of interest were the error rate, reaction time (RT), and lever-grip time. The error rate was calculated by dividing the total number of errors (early and later errors) by the total number of trials. RT was defined as the duration from a ‘go’ signal to the time point of lever release in a correct trial. Average error rate and RT were computed for each reward condition. The lever-grip time was defined as the duration from the end of ITI to the time when the monkey gripped the lever to initiate a trial (i.e., the latency to start a trial). Error rate and RT were analyzed using two-way repeated-measures ANOVAs with reward size (1, 2, 4, 8 drops) and satiation level (proportion of cumulative reward in a session: 0.125, 0.375, 0.625, 0.875) as within-subjects factors. The lever-grip time was analyzed using one-way repeated measures ANOVAs with satiation level (cumulative reward: 0.125, 0.375, 0.625, 0.875) as a within-subjects factor. The proportional behavioral data were transformed using the variance stabilizing arcsine transformation before hypothesis testing (Zar, 2013). The error rate was also analyzed by a model fitting as described previously (Minamimoto et al., 2009; Minamimoto et al., 2012). To assess the effects of reward size and satiation level on the error rate, the following model was used:

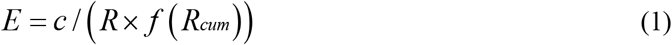

where *E* and *R* denote error rate and reward size, respectively. Parameter *c* is a monkey-specific parameter that represents reward-size sensitivity. *f(R_cum_)* denotes the reward discounting function, which was modeled as follows:

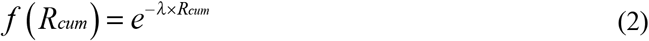

where *R_cum_* is a normalized cumulative reward in a session (0-1), and *λ* is a monkey-specific parameter that represents the steepness of reward discounting.

For neuronal data analysis, three task periods (cue period: 100-700 ms after cue on, pre-release period: 0-300 ms before lever release, reward period: 0-300 ms after reward delivery) and the baseline period (ITI: 0-500 ms before cue on) were defined. A neuron was classified as reward-size coding neuron when the firing rate during the task period was significantly modulated by reward size (main effect of reward size p < 0.05, one-way ANOVA) and linearly reflected reward size (p < 0.05, linear regression analysis). Neurons that showed positive or negative correlation were classified as positive or negative reward-size coding neurons, respectively. We also performed a non-parametric Kruskal-Wallis test to detect reward-size coding neurons (p < 0.05). The results based on the non-parametric analyses were not included if they confirmed the results from parametric analyses.

To quantify the time course of reward-size coding, the effect size (R squared) in a linear regression analysis with reward-size was calculated for every 100-ms window shift in 10-ms steps. Coding latency was defined as the duration between the cue onset and the time at which the first of three following consecutive 100-ms test intervals showed a significant reward-size effect (p < 0.05). Peak effect size was defined as the maximum effect size of individual neurons. Average coding latency and peak effect size were compared between VP and rmCD by Wilcoxon rank-sum test.

The effect of reward size and satiation level on firing rate during the task periods was also assessed by the following multiple linear regression model:

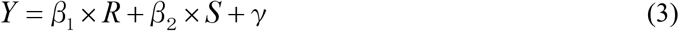

where *Y* is the firing rate, *R* is the reward size, *S* is the satiation level, *β_1_* and *β_2_* are the regression coefficients, and γ is a constant. Satiation level was inferred using equation (2) with the individual parameter *λ* derived from behavioral analysis. Neurons were classified into reward-size and satiation-level coding neurons if they had a significant correlation coefficient (p < 0.05) with each variable. A neuron was classified as a motivational-value coding neuron, when a neuron had a significant positive reward-size coefficient and negative satiation-level coefficient, or vice versa. The proportion of each type of neuron in the neuronal population (VP and rmCD) was calculated for each of the task periods, and compared between VP and rmCD using Wilcoxon rank-sum test with a threshold of statistical significance set by Bonferroni correction (alpha = 0.05/4). The proportion of motivational-value coding neurons was further compared to that of pseudo-motivational-value coding neurons, which were calculated by multiplying the proportion of neurons that coded satiation level and reward size orthogonally; this calculated the dual coding of two items of information by chance (i.e., joint probability).

For behavioral analysis of muscimol microinjection effects, the data obtained from two monkeys (monkeys RI and BI) were used for VP inactivation, while the data obtained from two monkeys (monkeys RI and RO) for rmCD inactivation (Nagai et al., 2016) were used for comparison. The dependent variables of interest were the change of error rate and early-error rate from the control session. The early-error rate was calculated by dividing the number of early error trials by that of total error trials (i.e., the sum of early and late error trials). RT and lever-grip time in the first and latter halves of sessions were also compared to assess the effects of satiation in control and muscimol sessions. Because VP inactivation induced similar effects in the two monkeys (monkeys BI and RI) regarding the elevation of error rate, the data were pooled across monkeys and compared to rmCD inactivation by Wilcoxon rank-sum test.

## Results

### Behavioral performance reflected reward size and satiation level

Two monkeys (TA and AP) learned to perform the reward-size task, in which unique visual cues provided information of the upcoming reward size (1, 2, 4, 8 drops of liquid; Fig. 1a and b). For both monkeys, error rate reflected reward size, such that the monkeys made more error responses (premature release or too-late release of the lever) when small rewards were assigned (Fig 1c). Error rate also reflected the satiation level, such that the error rate increased according to reward accumulation (Fig. 1d). Two-way repeated-measures ANOVAs (reward size: 1, 2, 4, 8 drops × cumulative reward: 0.125, 0.375, 0.625, 0.875) confirmed the significant effects of reward prediction and satiation on the behavioral performance (main effects of reward size and cumulative reward, and their interaction, p < 0.01, F > 11). As reported previously, the error rates were explained by a model in which the expected reward size was multiplied by an exponential decay function according to reward accumulation (Minamimoto et al., 2009; Minamimoto et al., 2012) (Fig. 1d, see Materials and Methods). The error rates were not influenced by reward history (p = 0.96, F_(1,2872)_ = 0.0, main effect of preceding reward size, three-way repeated measures ANOVAs), probably because reward size was randomly chosen and was independent from the previous trial.

The reaction time (RT) also changed in association with reward size and satiation level, such that the monkeys had a slower reaction for a smaller reward and when they had accumulated large amounts of rewards. Two-way repeated-measures ANOVAs revealed significant main effects of reward size and cumulative reward, and their interaction on RT (all p < 0.01, F > 3.5). The lever-grip time also changed according to the satiation level, such that the monkey tended to slowly grip the lever to initiate a trial in the later stage of a session. One-way repeated-measures ANOVAs revealed significant main effect of cumulative reward (p < 0.01, F_(1,178)_ = 102).

Together, these results suggest that the monkeys adjusted their motivation of the action based on the incentive value (i.e., expected reward size) and the internal drive (i.e., current satiation level).

### Task-related activity of VP and rmCD neurons

While monkeys TA and AP performed the reward-size task, we recorded the activity of 102 neurons (50 from TA and 52 from AP) in VP (Fig. 2a) and 106 neurons (68 from TA and 38 from AP) in rmCD (Fig. 2b). rmCD neurons were further classified into phasically-active neurons (PANs, n = 56), tonically-active neurons (TANs, n = 44) and fast-spiking neurons (FSNs, n = 6), based on criteria that were established in earlier studies (Aosaki et al., 1995; Yamada et al., 2016) (see Materials and Methods). The baseline firing rate (0-500 ms before cue, mean ± SEM) was high in VP (34.8 ± 1.8 spk/s) and low in rmCD (PANs 3.6 ± 0.4 spk/s, TANs 7.3 ± 0.4 spk/s, FSNs 14.3 ± 1.6 spk/s). In both areas, a majority of neurons showed excitatory response to the cue (VP: 44 excitatory and 16 inhibitory, rmCD: 39 excitatory and 17 inhibitory). Because the characteristics of neuronal activity among the three subtypes in rmCD were not significantly different in terms of value coding (i.e., reward-size and satiation-level coding) (p > 0.53, two-sample Kolmogorov-Smirnov test), we decided to treat them as a single population for the subsequent analyses. We will also report the results from PANs to ensure that the same conclusions would be reached.

### Neuronal activity in VP and rmCD reflected reward-size

We first examined how the incentive value is represented in VP and rmCD neurons. Fig. 3a-d illustrates four examples of neuronal activity showing reward-size modulation during the cue period. The first VP neuron example increased its activity after the largest reward (8 drops) cue, but decreased after smaller (1, 2, 4 drops) ones (Fig. 3a). The firing rate was positively correlated with the reward size (p < 0.01, r = 0.49, linear regression analysis, Fig. 3a, right), and therefore this neuron exhibited positive reward-size coding in this period. In another VP neuron example, the firing rate during the cue period became lower as a larger reward was expected (p < 0.01, r = −0.52, linear regression analysis), and thus this neuron exhibited negative reward-size coding (Fig. 3b). Similarly, we found that some rmCD neurons linearly encoded reward size during the cue period (Fig. 3c and d).

**Figure 3.**
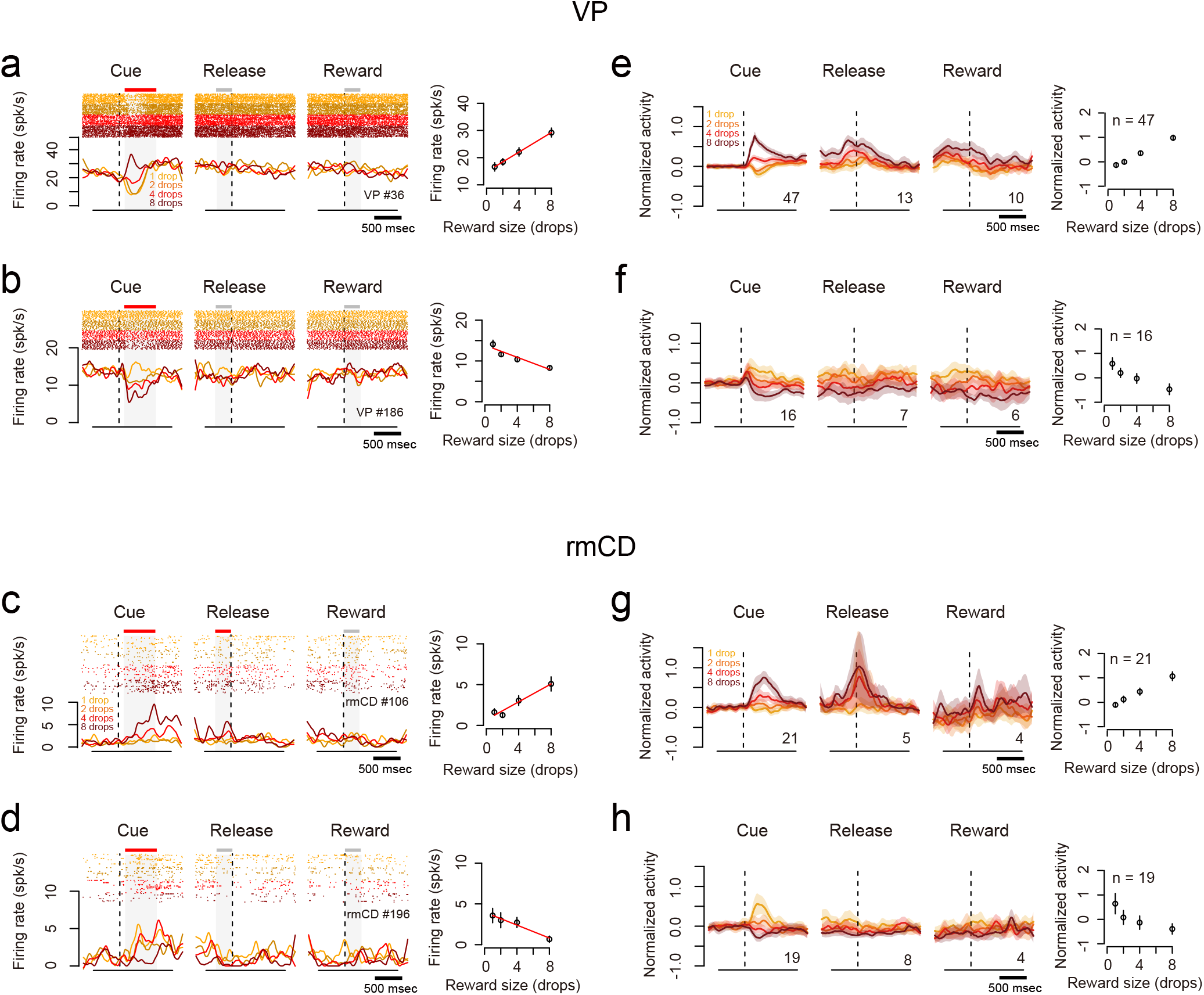
Reward-size coding in VP and rmCD. ***a-b***. Example activity of VP neurons showing positive (***a***) and negative (***b***) reward-size coding during cue period, respectively. Left: Raster spikes and spike density function (SDF, sigma = 10 ms) were aligned at task events. The colors correspond to the respective reward sizes. Red bars above shadings indicate significant linear correlation at the task period (p < 0.05, linear regression analysis), while gray bars indicate no significant correlation (p > 0.05). Right: Relationship between firing rate (mean ± SEM) during cue period and reward size. Regression lines are shown in red (p < 0.01). ***c-d.***. Examples of rmCD neurons. Schema of the figures are the same as in **a-b**. ***e-f***. Left: Population activities of VP neurons that were classified into positive (**e**) and negative (**f**) reward-size coding neurons, respectively. Curves and shades indicate mean and SEM of normalized activity to the baseline aligned at task events. Digits in each panel indicate the number of reward-size coding neurons at each task period. Right: Relationship between normalized neuronal activity during cue period (mean ± SEM) and reward size. ***g-h***. Population activities of rmCD neurons. Schema of the figures are the same as in **e-f**.

### VP had stronger reward-size coding than rmCD

We found many neurons that modulated their activity in at least one task period by the forthcoming reward size in both areas (VP: 71/102, rmCD: 51/106, p < 0.05, one-way ANOVA). Of these populations, most neurons linearly reflected reward size as we observed in Fig. 3 (VP: 68/71, rmCD: 46/51, p < 0.05, linear regression analysis). The proportion of reward-size coding neurons in VP was significantly larger than that in rmCD (p < 0.01, χ^2^ = 10, chi-square test). In rmCD, a large part of reward-size coding was observed in PANs (31/46). Reward-size coding was mainly observed during the cue period in both areas (VP 63/68, rmCD 40/46). As for population activity, both VP and rmCD clearly showed both types of linear reward-size coding during the cue period (Fig. 3e-h). During release or reward periods, however, linear reward-size coding was less clear.

To quantify reward-size coding, we computed the effect size (R squared) of activity in the sliding window (100 ms bin, 10 ms step) for each of the recorded neurons (Fig. 4a and b). Fig. 4 c and d show the average effect size of positive coding neurons (top) and negative coding neurons (bottom) in VP and rmCD, respectively. In both areas, the effect size rapidly and transiently increased after cue presentation, for both positive and negative coding neurons (Fig. 4c and d). The peak effect size of VP neurons (0.23 ± 0.016, median ± SEM) was significantly larger than that of rmCD neurons (0.14 ± 0.018) (p < 0.01, df = 101, rank-sum test) and that of PANs (0.14 ± 0.027, n = 26, p = 0.018, df = 87). Taken together, both VP and rmCD neurons exhibited reward-size modulation mainly after cue presentation, in which the former showed stronger modulation in terms of the proportion of neurons and effect size.

**Figure 4.**
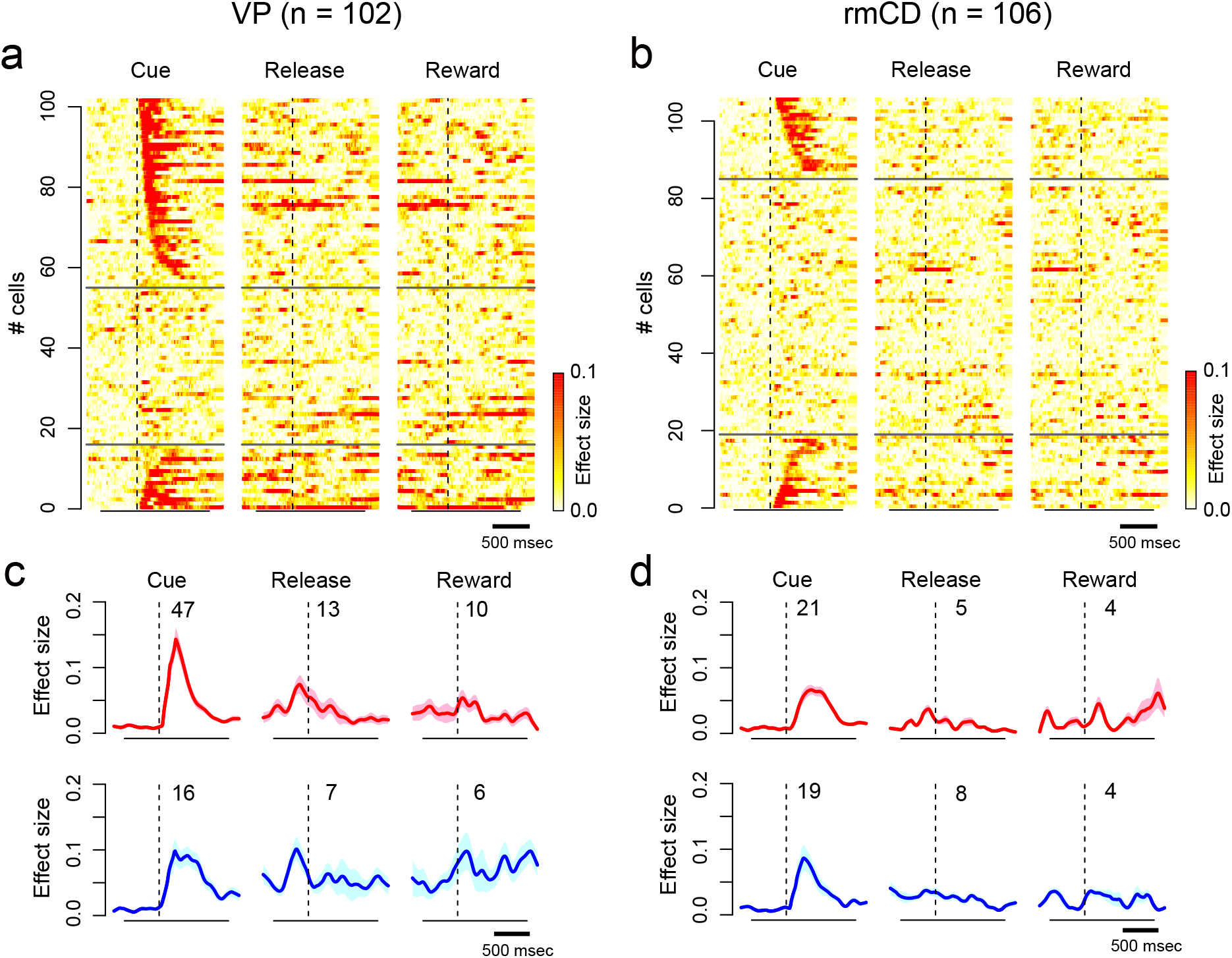
Time course of reward-size coding. ***a-b***. Time-dependent change of effect size (R squared) depicted with heat plots for VP neurons (***a***) and for rmCD neurons (***b***). Each panel shows data of each task event. Neurons are sorted by the coding latency from the cue. Upper rows show positive reward-size coding neurons, and lower rows show negative reward-size coding neurons. ***c-d***. Average effect size of positive (***top***) and negative (***bottom***) reward-size coding neurons around task events for VP neurons and for rmCD neurons. Digits in each panel indicate the number of reward-size coding neurons at each task period.

### Reward-size coding emerged earlier in VP than in rmCD

We compared the time course of reward-size coding after cue in two populations (VP, n = 63; rmCD, n = 40). The latency of reward-size coding of VP neurons was significantly shorter than that of rmCD neurons (VP 115 ± 17 ms, rmCD 225 ± 20 ms, median ± SEM; p < 0.01, df = 101, rank-sum test). The result held when we subsampled VP neurons with low baseline firing rate (< 25 spk/s, n = 32) and calculated the coding latency (120 ± 19 ms, p = 0.017, df = 70). Positive coding occurred earlier in VP than in rmCD (VP 100 ± 19 ms, rmCD 250 ± 27 ms, p < 0.01, df = 66, Fig. 5a and b), whereas the difference did not reach significance for negative coding (VP 150 ± 37 ms, rmCD 200 ± 29 ms, p = 0.48, df = 33, Fig. 5c and d).

**Figure 5.**
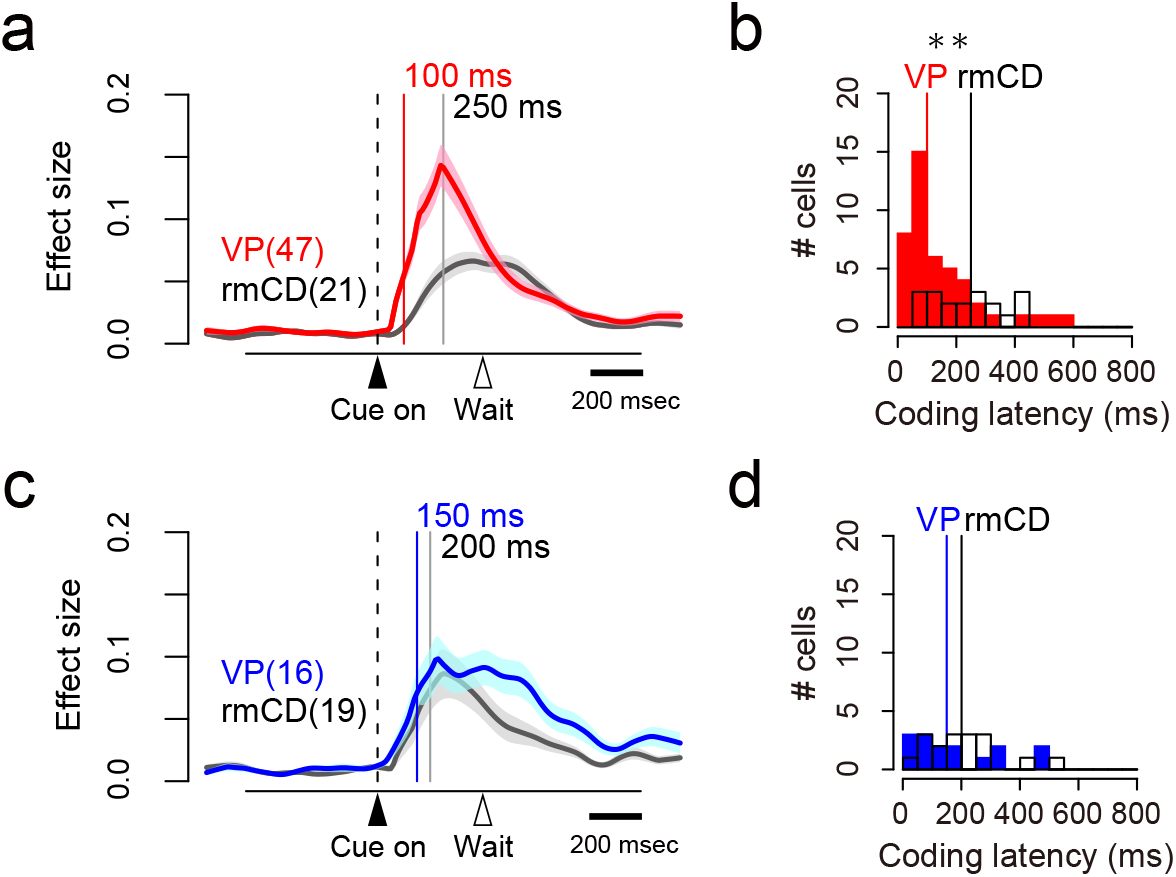
Coding latency of the expected reward size. ***a, c***. Effect size histogram aligned to cue for positive (***a***) and negative (***c***) reward-size coding neurons, reconstructed from Figure 4c-d. The data from VP (red and blue) and rmCD (gray) are depicted in the same panel. Vertical lines indicate the median of coding latency. ***b, d***. Distribution of coding latency for positive (***b***) and negative (***d***) reward-size coding neurons. Asterisk indicates significant difference between VP and rmCD (p < 0.01, rank-sum test).

If rmCD is the primary source for providing reward information to VP, the projection neurons (i.e., PANs) in rmCD would encode reward size earlier than VP neurons. However, the latency of reward-size coding of VP neurons was again shorter than that of PANs (VP 115 ± 17 ms, PANs 190 ± 25 ms, p = 0.049, df = 87).

### Encoding of satiation level in VP and rmCD

As shown above, the monkeys’ goal-directed behavior (i.e., error rate) is affected by internal drive (i.e., satiation change) as well as incentive value (Fig. 1d). We found that some neurons changed their firing rate according to the satiation level. For instance, a VP neuron decreased its activity after the cue with reward-size (negative reward-size coding), while the decrease became smaller according to reward accumulation (Fig. 6a-c). This was not due to changes in isolation during a recording session as confirmed by unchanged spike waveforms (Fig. 6d). In an example of an rmCD neuron, the firing rate in the cue period was positively related to reward size and negatively related to reward accumulation (Fig. 6e-h).

**Figure 6.**
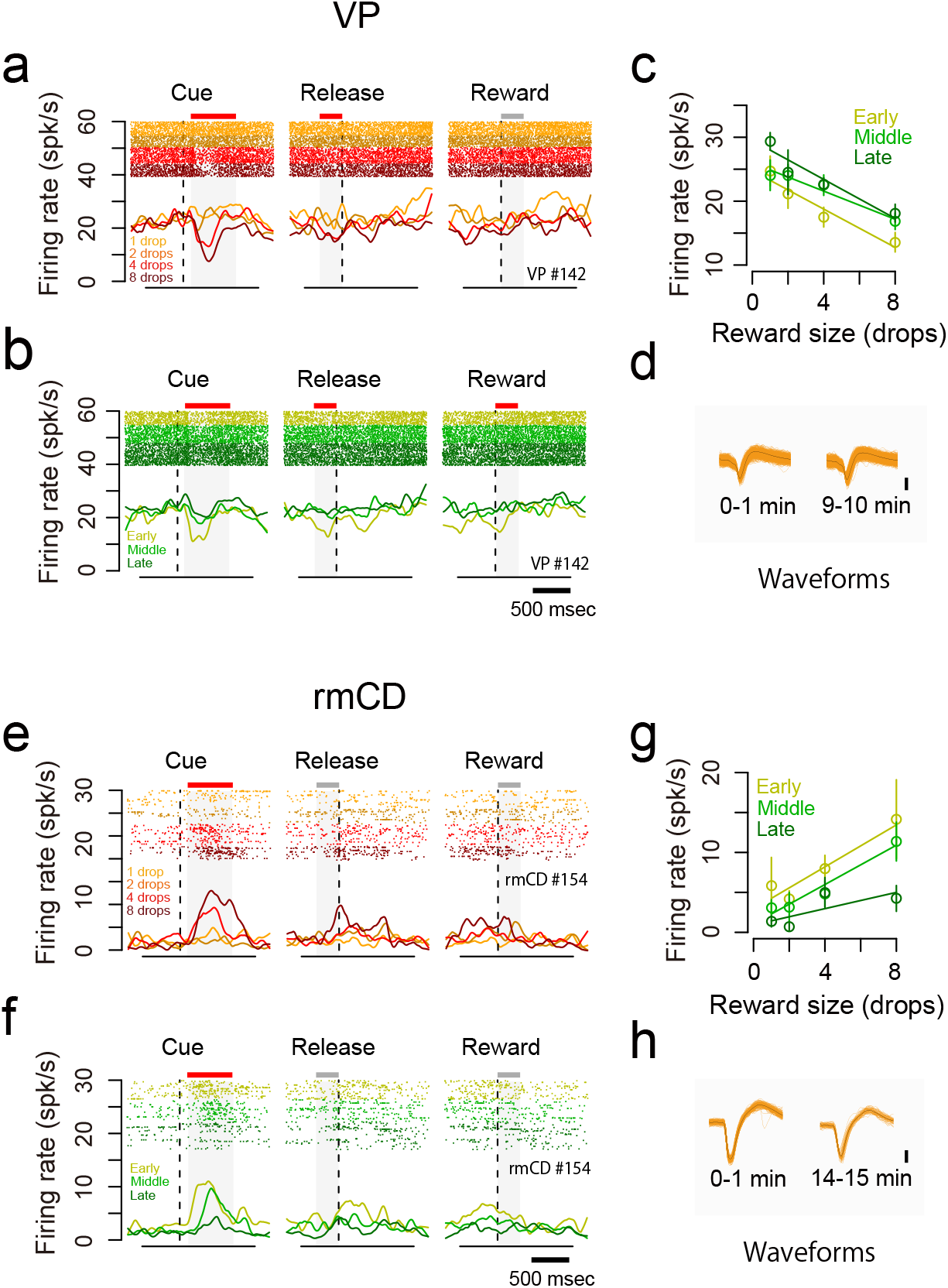
Dual coding of reward size and satiation level in single neurons. ***a-d***. An example VP neuron showing negative reward-size coding and positive satiation-level coding. ***a-b***. Raster plots and SDF are shown for each reward condition (***a***) and for each session period (***b***). ***c***. The relationship between firing rate during the cue period (mean ± SEM) and reward size are plotted for the session period. Linear regressions are shown as colored lines. ***d***. Waveforms of each spike (orange) and average waveform (black) during the first minute (left) and last minute (right) in the recording session. ***e-h***. An example rmCD neuron showing positive reward-size coding and negative satiation-level coding. Schema of the figures are the same as in ***a-d***.

To assess how satiation level and reward size were encoded in VP and rmCD, we performed a multiple linear regression analysis on the activity during each of four task periods (ITI, cue, pre-release, reward). For this analysis, the satiation level was inferred using the model with monkey-specific parameter *λ* that explained individual behavioral data (Fig. 1d, see Materials and Methods). Fig. 7a shows scatter plots of standardized regression coefficient for satiation level and reward size for the activity during the cue period of individual neurons in VP and rmCD. The proportion of satiation-level coding neurons during the cue period was not significantly different between the two areas (p = 0.28, χ^2^ = 1.2, chi-square test). When we ran the same analysis for other periods (Fig. 7b left), this proportion was significantly larger in VP than in rmCD (p < 0.05 with Bonferroni correction, rank-sum test). In contrast, the proportion of reward-size coding neurons in VP was larger than that in rmCD for the cue period (p < 0.01, χ^2^ = 11), but not for the other task periods (p > 0.05 with Bonferroni correction) (Fig. 7b center). A similar tendency was found when we compared VP and PANs; satiation-level coding was more frequent in VP than in PANs during the ITI, pre-release, and reward periods (p < 0.05), while reward-size coding tended to be more frequent in VP than in PANs during the cue period (p = 0.10, χ^2^ = 2.7).

**Figure 7.**
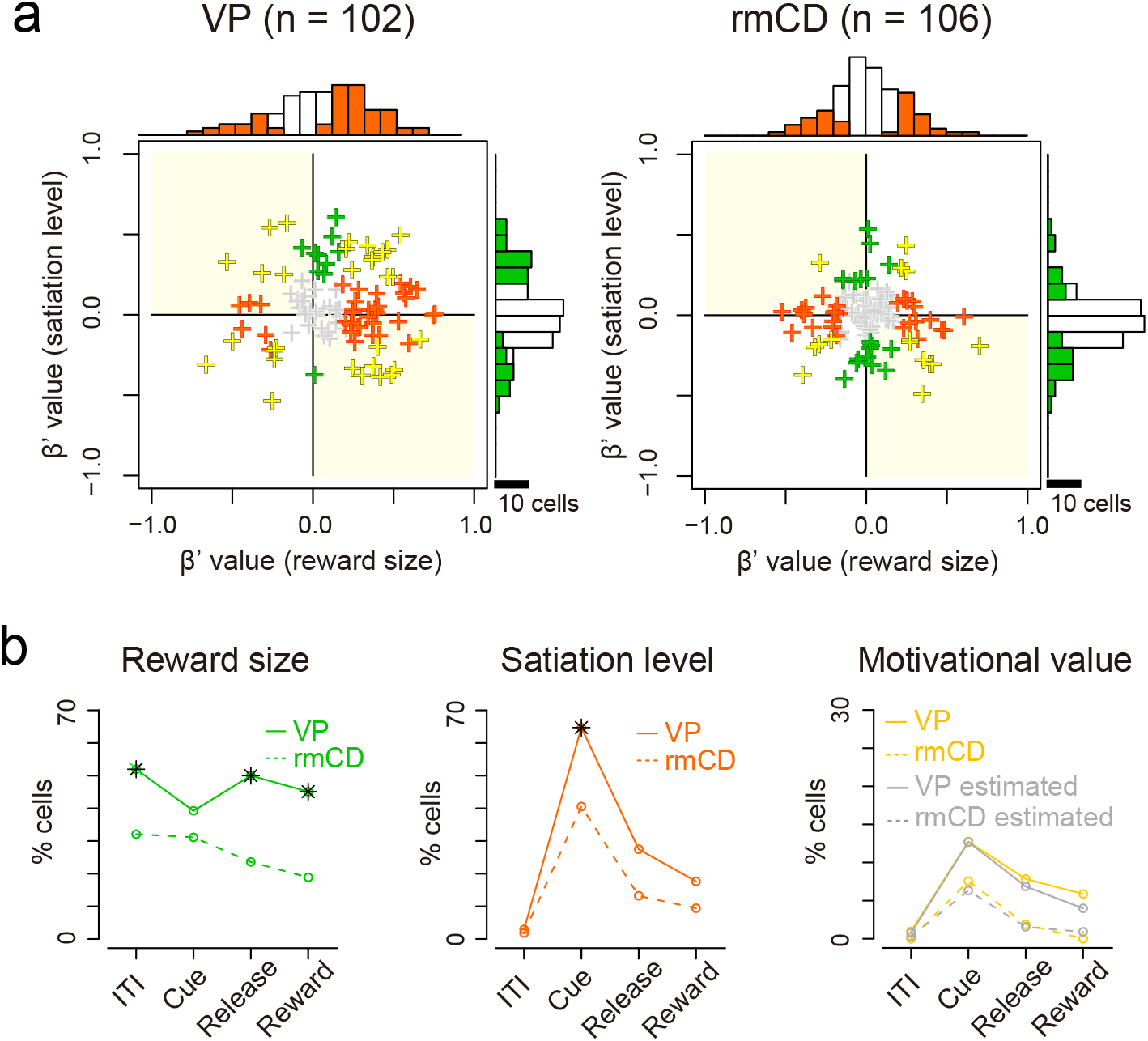
Separate coding of reward-size and satiation-level in VP and rmCD. ***a***. Scatter plot of standardized correlation coefficients (cue period) for reward size (abscissa) against satiation level (ordinate) are shown for VP neurons (***left***) and rmCD neurons (***right***), respectively. Colors indicate significant reward-size coding neurons (orange), satiation-level coding neurons (green), motivational-value coding neurons (yellow), and non-coding neurons (gray). Histograms in the main panels illustrate the distribution of coefficients with significant neurons with satiation level and reward size. ***b***. Proportion of VP neurons (solid line) and rmCD neurons (dashed line) that showed satiation-level coding (***left***), reward-size coding (***center***) and motivational-value coding (***right***) for each task period (ITI, cue, pre-release, and reward periods). Asterisks indicate significant difference between VP and rmCD (p < 0.05 with Bonferroni correction, chi-square test). For motivational-value coding (***right***), proportion of neurons with pseudo-motivational value coding by chance is shown in gray. No significant difference was observed between data and estimation (p > 0.05 with Bonferroni correction, chi-square test).

By definition, motivational value increases as expected reward size increases, and as satiation level decreases (Berridge, 2004; Zhang et al., 2009). Thus, motivational value coding neuron is defined as a neuron that showed positive reward-size coding and negative satiation-level coding, or vice versa (Fig. 7a, yellow areas; see Fig. 6 for examples). We found some motivational-value coding neurons during cue, pre-release and reward periods in both areas (Fig. 7b right). However, the proportion was not significantly larger than that of neurons by chance coding both satiation level and reward size in the opposite direction (all P > 0.05, chi-square test), suggesting independent signaling of reward size and satiation level in VP and rmCD.

In our paradigm, the satiation level monotonically changed according to reward accumulation and time spent during a session, and thus both are highly correlated. To examine the possibility of time elapsed rather than reward accumulation accounting for satiation coding, we divided the neurons into “rapid-half” and “slow-half” based on the reward accumulation speed of recording session (drops/min) in which the neuron was recorded. We found that the “rapid-half” group contained more satiation-level coding neurons (rapid: 24, slow: 16) and showed stronger effect size (beta value) on average (rapid: 0.20, slow: 0.17) in VP, confirming a larger contribution of reward accumulation to neuronal modulation.

### Inactivation of VP disrupted goal-directed behavior

Previous study demonstrated that bilateral inactivation of rmCD by local injection of the GABA_A_ receptor agonist muscimol diminished reward-size sensitivity (Nagai et al., 2016) as indicated by an increase in the error rate of reward-size task especially in larger reward-size trials (Fig. 8a), supporting that the neuronal activity in rmCD is essential for controlling goal-directed behavior based on the expected reward size.

**Figure 8.**
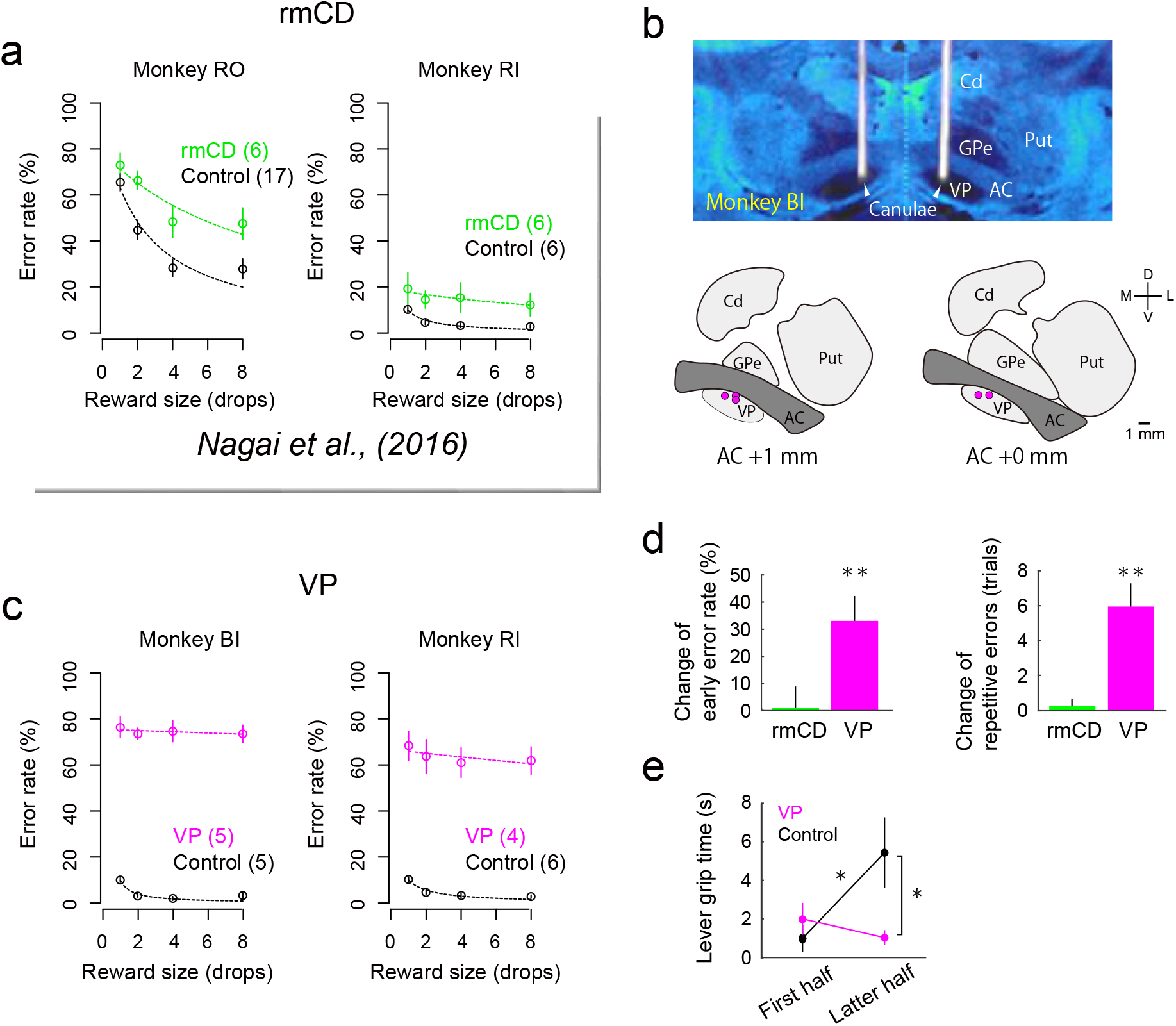
Behavioral change due to inactivation of VP and rmCD. ***a***. Error rate in control (black) and rmCD inactivation (green) sessions for monkey RO (***left***) and monkey RI (***right***). Digits in panels indicate number of sessions. ***b***. Injection sites in VP. Top panel shows representative CT/MR fusion image for confirmation of injection sites in monkey BI. Bottom two panels illustrate the location of injection indicated by magenta dots. ***c***. Error rate in control (black) and VP inactivation (magenta) sessions for monkeys BI (***left***) and RI (***right***), respectively. ***d***. Change of early-error rate (***left***) and length of repetitive errors (***right***) by rmCD inactivation (green) and VP inactivation (magenta). Asterisks indicate significant differences from control session (* p < 0.05, ** p < 0.01, rank-sum test). ***e***. Lever-grip time in the first and latter half of control session (black) and of VP inactivation session (magenta) (mean ± SEM). Asterisks indicate significant differences by post-hoc Tukey test (* p < 0.05).

To examine how VP contributes to the control of goal-directed behavior, we injected muscimol into bilateral VP (3 μg/μL, 2 μL/side) of two monkeys (monkeys BI and RI) and tested them with reward-size task. CT images visualizing the injection cannulae confirmed the sites of muscimol injection; they were located in the VP matching the recording sites (Fig. 8b; see Fig. 2a for comparison). Bilateral VP inactivation significantly increased the error rate regardless of reward size (main effect of treatment, p < 0.01, F_(1, 17)_ = 449.7, repeated measures ANOVAs, Fig. 8c). The increase in error rate by VP inactivation was significantly greater than that by rmCD inactivation (data pooled across monkeys, p < 0.01, df = 18, rank-sum test). VP inactivation did not decrease, but rather increased the total number of trials performed (control 922 ± 132 trials; VP inactivation 1464 ± 153 trials; mean ± SEM). VP inactivation did not change RT overall (main effect of treatment, p = 0.15, F_(1, 17)_ = 2.2, repeated measures ANOVAs). After VP inactivation, the monkeys frequently released the lever before the go signal appeared, or even before red or cue came on, and thereby increased the proportion of early errors (p < 0.01, df = 16, rank-sum test; Fig. 8d left) and error repetition (p < 0.01, df = 16, Fig. 8d right). The error pattern changes were not observed after rmCD inactivation (p > 0.47, df = 22, Fig. 8d).

VP inactivation also totally abolished the relationship among error rates, reward size and satiation; the error rate was not explained by the model used in the control condition (Eq. 1; R^2^ = 0.0, cf. Fig. 1d). While the lever-grip time was extended in the latter half of a session in the control session, it was insensitive to satiation after VP inactivation, resulting in a significant interaction (satiation × treatment) by two-way ANOVA (p = 0.025, F_(1,36)_ = 5.5, Fig. 8e). VP inactivation did not affect the satiation effect on RT (two-way ANOVA, interaction, p = 0.52, F_(1,36)_ = 0.43; main effect of satiation, p = 0.018, F_(1,36)_ = 6.1). Taken together, these results suggest that inactivation of VP disrupted normal goal-directed control of behavior including loss of reward-size and satiation effects.

## Discussion

In the present study, we examined the activity of VP and rmCD neurons during goal-directed behavior controlled by both incentive value (i.e., reward size) and internal drive (i.e., satiation level). We found that reward-size coding after a reward-size cue was stronger and earlier in VP neurons than in rmCD neurons. We also found that satiation-level coding was observed throughout a trial, and appeared more frequently in VP than in rmCD neurons. In both areas, information regarding reward size and satiation level was not systematically integrated into a single neuron but was independently signaled in the population. Inactivation of the bilateral VP disrupted normal goal-directed control of action, suggesting a causal role of VP in signaling incentive and drive for motivational control of goal-directed behavior.

Past studies demonstrated that neurons in VP and rmCD encode the incentive value of cue during performing the task offering binary outcomes (e.g., large or small reward) (Tindell et al., 2004; Nakamura et al., 2012; Tachibana and Hikosaka, 2012; Saga et al., 2017). Instead, the reward-size task used in the present study offered four reward sizes that enabled us to quantify the linearity of value coding in the activity of single neurons. With this paradigm, we found that some rmCD neurons exhibited the activity reflecting reward-size mainly during cue periods. The activity is likely to mediate goal-directed behavior based on expected reward size, because inactivation of this brain area impaired the task performance by the loss of reward-size sensitivity (Nagai et al., 2016). Our data are also consistent with previous research, which proposed the projection from the OFC to the striatum as being the critical pathway of carrying incentive information and performing a goal-directed behavior (Yin et al., 2005; Gremel and Costa, 2013; Gremel et al., 2016); we thereby confirmed the role of rmCD for mediating goal-directed behavior.

The present results, however, highlighted the more prominent role of VP in signaling incentive information for goal-directed behavior. We found that neuronal modulation by expected value was stronger and more frequent in VP neurons than in rmCD neurons. The strong signal in VP could not be accounted for solely by the convergence input from the rmCD for two reasons. First, the coding latency of VP neurons was significantly shorter than that of rmCD projection neurons (PANs). Second, if this were the case, response polarity would be opposite between the two areas, considering that the projection from rmCD to VP is GABAergic (Haber et al., 1990). However, a majority of the neurons in both areas showed excitatory response to cue. One might argue that the reward-size coding in rmCD neurons would be non-monotonic (e.g., neurons exclusively respond to a certain reward size) and thereby the coding strength of rmCD would have been underestimated. This was not likely, however, because we did not find such neurons when we checked the SDF of each neuron, and because we observed only a few neurons that showed significance by one-way ANOVA but not by linear regression. Together, our results suggest that VP signals incentive value that does not primarily originate from rmCD. This suggestion may also extend to the limbic striatum, given the previous finding of similar earlier incentive signaling in VP than in the nucleus accumbens in rats (Richard et al., 2016).

A remaining question is: where is such rich and rapid incentive-value information derived from? One possible source is the basolateral amygdala (BLA), which has a reciprocal connection to VP (Mitrovic and Napier, 1998; Root et al., 2015) and is known to contain neurons reflecting incentive value of cue with short latency (Paton et al., 2006; Belova et al., 2008; Jenison et al., 2011). Recent studies demonstrated that amygdala lesion impaired reward-based learning more severely than VS lesion in monkeys (Averbeck et al., 2014; Costa et al., 2016), supporting the contribution of BLA-VP projection in incentive-value processing. Another candidate is the projection from the subthalamic nucleus (STN); VP has a reciprocal connection with the medial STN that receives projections from limbic cortical areas (Haynes and Haber, 2013), composing the limbic cortico-subthalamo-pallidal ‘hyperdirect’ pathway (Nambu et al., 2002). It has been shown that STN neurons respond to cues predicting rewards in monkeys (Matsumura et al., 1992; Darbaky et al., 2005; Espinosa-Parrilla et al., 2015). Future studies should identify the source of the incentive-value information in terms of the latency, strength and linearity of the coding.

In addition to reward-size coding, VP neurons also encoded the internal drive (satiation level) of monkeys, although other factors (e.g., time elapsed, fatigue, etc.) could be confounded. This satiation-level coding was prominent even in the ITI phase, suggesting that this is indeed a reflection of motivational state rather than task structure *per se*. A similar type of state coding has been reported in agouti-related peptide (AgRP) producing neurons in the arcuate nucleus of the hypothalamus (ARH); hunger/satiety state modulates the firing rate of ARH-AgRP neurons (Chen et al., 2015), which regulates feeding behavior together with the lateral hypothalamus (LH) (Burton et al., 1976; Petrovich, 2018). Given that VP receives direct input from the hypothalamus (Castro et al., 2015; Root et al., 2015), the satiation coding in VP might reflect the state-dependent activity originating from the hypothalamus. Although the current results indicate that both incentive value and internal drive are not systematically integrated into a single neuron level, VP may play a pivotal role in representing the two factors, which may be integrated in downstream structures, such as the mediodorsal (MD) thalamus (Haber and Knutson, 2010). MD is one of the brain regions responsible for ‘reinforcer devaluation’, i.e., appropriate action selection according to the satiation of specific needs (Mitchell et al., 2007; Izquierdo and Murray, 2010), and therefore motivational value could be formulated in this area.

By definition, the motivational value refers the value that is directly linked to goal-directed action. A similar value system, the subjective value that is inferred from choice behavior, also regards the subject’s internal state (Bernoulli, 1982; Stephens and Krebs, 1986). A crucial role for subject-centered decision has been demonstrated in the ventromedial prefrontal cortex (vmPFC), where neural activity encodes subjective value (de Araujo et al., 2003; Kable and Glimcher, 2007; Bouret and Richmond, 2010; O’Doherty, 2011). Because vmPFC is one of the projection targets from MD (Goldman-Rakic and Porrino, 1985; McFarland and Haber, 2002), its value coding could be indirectly modulated by signal originating from VP. Therefore, the current results may provide a subcortical mechanism of value coding that contributes to value-based decision-making as well as goal-directed control of action.

The causal contribution of value coding in VP was examined by an inactivation study. We found that bilateral inactivation of VP increased premature errors irrespective of incentive conditions. VP inactivation had little effect on the general sensory-motor process, motivation or arousal, because the inactivation did not change RT or decrease the number of trials performed. VP inactivation attenuated the satiation effect on lever-grip time, which is considered to reflect the motivational state (Kobayashi et al., 2002). Failure of model fitting after VP inactivation also suggests impairment of motivational control based on incentive and drive. The present results together with a previous study support the view that the value coding of VP contributes to the motivational control of goal-directed behavior (Tachibana and Hikosaka, 2012). Another mechanism is also possible, such as, that suppressing general high neuronal activity in VP (cf. Fig. 3) would promote a premature response. Inactivation of VP would activate its efferent target neurons including dopamine neurons by a disinhibition mechanism, and thereby abnormally invigorate current actions (Niv et al., 2007; Tachibana and Hikosaka, 2012). However, this story is not so simple, as injection of the GABA_A_ receptor antagonist bicuculline into bilateral VP also increased premature responses in monkeys (Saga et al., 2017). Thus, disruption of value coding in VP may be the fundamental mechanism underlying the observed abnormal behavior. The loss of information regarding motivational value could promote a shift from goal-directed to habitual control of action (Dickinson, 1985; Dickinson and Balleine, 1994), which is implicated in the hallmark of addictive disorders (Everitt and Robbins, 2005; Ersche et al., 2016). Our results, therefore, emphasize the importance of future investigations into the exact neuronal mechanisms of motivational value formulation and control of actions in both the normal and abnormal state.

In conclusion, our data highlight the critical contribution of VP in goal-directed action. VP neurons independently encode information regarding incentive and drive that are essential for motivational control of goal-directed behavior. Regarding this view, VP may gain access to motor-related processes and adjust the motivation of action based on the expected reward value in accordance with the current needs.

## Acknowledgments

We thank J. Kamei, Y. Matsuda, R. Yamaguchi, Y. Sugii and R. Suma for their technical assistance, and Dr. I. Monosov for his invaluable technical advice and discussion. This work was supported by JSPS KAKENHI [JP15H05917] (to T. M.), [JP15H06872, JP17K13275] (to A. F.) from the Ministry of Education, Culture, Sports, Science, and Technology of Japan (MEXT), and by the Strategic Research Program for Brain Sciences from the Japan Agency for Medical Research and Development (AMED) JP18dm0107146 (to T. M.).

Author Contributions
A. F. and T. M. designed the research; A. F., Y. H., Y. N. and E. K. performed the research; A. F. analyzed the data; all authors wrote the manuscript.

## References

Ahrens AM, Meyer PJ, Ferguson LM, Robinson TE, Aldridge JW (2016) Neural Activity in the Ventral Pallidum Encodes Variation in the Incentive Value of a Reward Cue. J Neurosci 36:7957–7970.

Aosaki T, Kimura M, Graybiel AM (1995) Temporal and spatial characteristics of tonically active neurons of the primate’s striatum. J Neurophysiol 73:1234–1252.

Averbeck BB, Lehman J, Jacobson M, Haber SN (2014) Estimates of projection overlap and zones of convergence within frontal-striatal circuits. J Neurosci 34:9497–9505.

Belova MA, Paton JJ, Salzman CD (2008) Moment-to-moment tracking of state value in the amygdala. J Neurosci 28:10023–10030.

Bernoulli D (1982) The works. In: Birkhauser, Boston.

Berridge KC (2004) Motivation concepts in behavioral neuroscience. Physiol Behav 81:179–209.

Bouret S, Richmond BJ (2010) Ventromedial and orbital prefrontal neurons differentially encode internally and externally driven motivational values in monkeys. J Neurosci 30:8591–8601.

Burton MJ, Rolls ET, Mora F (1976) Effects of hunger on the responses of neurons in the lateral hypothalamus to the sight and taste of food. Exp Neurol 51:668–677.

Castro DC, Cole SL, Berridge KC (2015) Lateral hypothalamus, nucleus accumbens, and ventral pallidum roles in eating and hunger: interactions between homeostatic and reward circuitry. Front Syst Neurosci 9:90.

Chen Y, Lin YC, Kuo TW, Knight ZA (2015) Sensory detection of food rapidly modulates arcuate feeding circuits. Cell 160:829–841.

Costa VD, Dal Monte O, Lucas DR, Murray EA, Averbeck BB (2016) Amygdala and Ventral Striatum Make Distinct Contributions to Reinforcement Learning. Neuron 92:505–517.

Darbaky Y, Baunez C, Arecchi P, Legallet E, Apicella P (2005) Reward-related neuronal activity in the subthalamic nucleus of the monkey. Neuroreport 16:1241–1244.

de Araujo IE, Kringelbach ML, Rolls ET, McGlone F (2003) Human cortical responses to water in the mouth, and the effects of thirst. J Neurophysiol 90:1865–1876.

Dickinson A (1985) Actions and habits: the development of behavioural autonomy. Phil Trans R Soc Lond B 308:67–78.

Dickinson A, Balleine B (1994) Motivational control of goal-directed action. Animal Learning & Behavior 22:1–18.

Ersche KD, Gillan CM, Jones PS, Williams GB, Ward LH, Luijten M, de Wit S, Sahakian BJ, Bullmore ET, Robbins TW (2016) Carrots and sticks fail to change behavior in cocaine addiction. Science 352:1468–1471.

Espinosa-Parrilla JF, Baunez C, Apicella P (2015) Modulation of neuronal activity by reward identity in the monkey subthalamic nucleus. Eur J Neurosci 42:1705–1717.

Everitt BJ, Robbins TW (2005) Neural systems of reinforcement for drug addiction: from actions to habits to compulsion. Nat Neurosci 8:1481–1489.

Faget L, Zell V, Souter E, McPherson A, Ressler R, Gutierrez-Reed N, Yoo JH, Dulcis D, Hnasko TS (2018) Opponent control of behavioral reinforcement by inhibitory and excitatory projections from the ventral pallidum. Nat Commun 9:849.

Goldman-Rakic PS, Porrino LJ (1985) The primate mediodorsal (MD) nucleus and its projection to the frontal lobe. J Comp Neurol 242:535–560.

Gremel CM, Costa RM (2013) Orbitofrontal and striatal circuits dynamically encode the shift between goal-directed and habitual actions. Nat Commun 4:2264.

Gremel CM, Chancey JH, Atwood BK, Luo G, Neve R, Ramakrishnan C, Deisseroth K, Lovinger DM, Costa RM (2016) Endocannabinoid Modulation of Orbitostriatal Circuits Gates Habit Formation. Neuron 90:1312–1324.

Groenewegen HJ, Berendse HW, Haber SN (1993) Organization of the output of the ventral striatopallidal system in the rat: ventral pallidal efferents. Neuroscience 57:113–142.

Haber SN, Knutson B (2010) The reward circuit: linking primate anatomy and human imaging. Neuropsychopharmacology 35:4–26.

Haber SN, Groenewegen HJ, Grove EA, Nauta WJ (1985) Efferent connections of the ventral pallidum: evidence of a dual striato pallidofugal pathway. J Comp Neurol 235:322–335.

Haber SN, Lynd E, Klein C, Groenewegen HJ (1990) Topographic organization of the ventral striatal efferent projections in the rhesus monkey: an anterograde tracing study. J Comp Neurol 293:282–298.

Haynes WI, Haber SN (2013) The organization of prefrontal-subthalamic inputs in primates provides an anatomical substrate for both functional specificity and integration: implications for Basal Ganglia models and deep brain stimulation. J Neurosci 33:4804–4814.

Hays Jr A, Richmond B, Optican L (1982) Unix-based multiple-process system, for real-time data acquisition and control.

Izquierdo A, Murray EA (2010) Functional interaction of medial mediodorsal thalamic nucleus but not nucleus accumbens with amygdala and orbital prefrontal cortex is essential for adaptive response selection after reinforcer devaluation. J Neurosci 30:661–669.

Jenison RL, Rangel A, Oya H, Kawasaki H, Howard MA (2011) Value encoding in single neurons in the human amygdala during decision making. J Neurosci 31:331–338.

Kable JW, Glimcher PW (2007) The neural correlates of subjective value during intertemporal choice. Nat Neurosci 10:1625–1633.

Kobayashi Y, Inoue Y, Yamamoto M, Isa T, Aizawa H (2002) Contribution of pedunculopontine tegmental nucleus neurons to performance of visually guided saccade tasks in monkeys. J Neurophysiol 88:715–731.

Mahler SV, Vazey EM, Beckley JT, Keistler CR, McGlinchey EM, Kaufling J, Wilson SP, Deisseroth K, Woodward JJ, Aston-Jones G (2014) Designer receptors show role for ventral pallidum input to ventral tegmental area in cocaine seeking. Nat Neurosci 17:577–585.

Mai JK, Paxinos G (2011) The human nervous system: Academic Press.

Matsumura M, Kojima J, Gardiner TW, Hikosaka O (1992) Visual and oculomotor functions of monkey subthalamic nucleus. J Neurophysiol 67:1615–1632.

McFarland NR, Haber SN (2002) Thalamic relay nuclei of the basal ganglia form both reciprocal and nonreciprocal cortical connections, linking multiple frontal cortical areas. J Neurosci 22:8117–8132.

Minamimoto T, La Camera G, Richmond BJ (2009) Measuring and modeling the interaction among reward size, delay to reward, and satiation level on motivation in monkeys. J Neurophysiol 101:437–447.

Minamimoto T, Hori Y, Richmond BJ (2012) Is working more costly than waiting in monkeys? PLoS One 7:e48434.

Mitchell AS, Browning PG, Baxter MG (2007) Neurotoxic lesions of the medial mediodorsal nucleus of the thalamus disrupt reinforcer devaluation effects in rhesus monkeys. J Neurosci 27:11289–11295.

Mitrovic I, Napier TC (1998) Substance P attenuates and DAMGO potentiates amygdala glutamatergic neurotransmission within the ventral pallidum. Brain Res 792:193–206.

Nagai Y et al. (2016) PET imaging-guided chemogenetic silencing reveals a critical role of primate rostromedial caudate in reward evaluation. Nat Commun 7:13605.

Nakamura K, Santos GS, Matsuzaki R, Nakahara H (2012) Differential reward coding in the subdivisions of the primate caudate during an oculomotor task. J Neurosci 32:15963–15982.

Nambu A, Tokuno H, Takada M (2002) Functional significance of the cortico-subthalamo-pallidal ‘hyperdirect’ pathway. Neurosci Res 43:111–117.

Niv Y, Daw ND, Joel D, Dayan P (2007) Tonic dopamine: opportunity costs and the control of response vigor. Psychopharmacology (Berl) 191:507–520.

O’Doherty JP (2011) Contributions of the ventromedial prefrontal cortex to goal-directed action selection. Ann N Y Acad Sci 1239:118–129.

Paton JJ, Belova MA, Morrison SE, Salzman CD (2006) The primate amygdala represents the positive and negative value of visual stimuli during learning. Nature 439:865–870.

Petrovich GD (2018) Lateral Hypothalamus as a Motivation-Cognition Interface in the Control of Feeding Behavior. Front Syst Neurosci 12:14.

Ray JP, Price JL (1993) The organization of projections from the mediodorsal nucleus of the thalamus to orbital and medial prefrontal cortex in macaque monkeys. J Comp Neurol 337:1–31.

Richard JM, Ambroggi F, Janak PH, Fields HL (2016) Ventral Pallidum Neurons Encode Incentive Value and Promote Cue-Elicited Instrumental Actions. Neuron 90:1165–1173.

Rolls ET (2006) Brain mechanisms underlying flavour and appetite. Philos Trans R Soc Lond B Biol Sci 361:1123–1136.

Root DH, Melendez RI, Zaborszky L, Napier TC (2015) The ventral pallidum: Subregion-specific functional anatomy and roles in motivated behaviors. Prog Neurobiol 130:29–70.

Saga Y, Richard A, Sgambato-Faure V, Hoshi E, Tobler PN, Tremblay L (2017) Ventral Pallidum Encodes Contextual Information and Controls Aversive Behaviors. Cereb Cortex 27:2528–2543.

Smith KS, Tindell AJ, Aldridge JW, Berridge KC (2009) Ventral pallidum roles in reward and motivation. Behav Brain Res 196:155–167.

Stephens DW, Krebs JR (1986) Foraging theory: Princeton University Press.

Tachibana Y, Hikosaka O (2012) The primate ventral pallidum encodes expected reward value and regulates motor action. Neuron 76:826–837.

Tindell AJ, Berridge KC, Aldridge JW (2004) Ventral pallidal representation of pavlovian cues and reward: population and rate codes. J Neurosci 24:1058–1069.

Yamada H, Inokawa H, Hori Y, Pan X, Matsuzaki R, Nakamura K, Samejima K, Shidara M, Kimura M, Sakagami M, Minamimoto T (2016) Characteristics of fast-spiking neurons in the striatum of behaving monkeys. Neurosci Res 105:2–18.

Yin HH, Knowlton BJ, Balleine BW (2005) Blockade of NMDA receptors in the dorsomedial striatum prevents action-outcome learning in instrumental conditioning. Eur J Neurosci 22:505–512.

Zar JH (2013) Biostatistical Analysis: Pearson New International Edition: Pearson Higher Ed.

Zhang J, Berridge KC, Tindell AJ, Smith KS, Aldridge JW (2009) A neural computational model of incentive salience. PLoS Comput Biol 5:e1000437.

